# Combining Inferred Regulatory and Reconstructed Metabolic Networks Enhances Phenotype Prediction in Yeast

**DOI:** 10.1101/087148

**Authors:** Zhuo Wang, Samuel A. Danziger, Benjamin D. Heavner, Shuyi Ma, Jennifer J. Smith, Song Li, Thurston Herricks, Evangelos Simeonidis, Nitin S. Baliga, John D. Aitchison, Nathan D. Price

**Affiliations:** The Bio-X Institute, School of Life Sciences and Biotechnology, Shanghai Jiao Tong University, Shanghai, 200240, China; Institute for Systems Biology, Seattle, WA, 98109, USA; Center for Infectious Disease Research, Seattle, WA, 98109, USA; Department of Biostatistics, University of Washington, Seattle, WA, 98195, USA; Department of Chemical and Biomolecular Engineering, University of Illinois, Urbana-Champaign, IL, 61801, USA; Departments of Biology and Microbiology & Molecular and Cellular Biology Program, University of Washington, Seattle, WA, 98195, USA; Lawrence Berkeley National Lab, Berkeley, CA, 94720, USA

**Author notes:** **Correspondence**: **Nitin S. Baliga**, **John D. Aitchison**, **Nathan D. Price**.

**Keywords:** metabolic network, Environment and Gene Regulatory Influence Network (EGRIN), Probabilistic Regulation of Metabolism (PROM), metabolic flux, synthetic lethal

## Abstract

Gene regulatory and metabolic network models have been used successfully in many organisms, but inherent differences between them make networks difficult to integrate. Probabilistic Regulation Of Metabolism (PROM) provides a partial solution, but it does not incorporate network inference and underperforms in eukaryotes. We present an Integrated Deduced REgulation And Metabolism (IDREAM) method that combines statistically inferred Environment and Gene Regulatory Influence Network (EGRIN) models with the PROM framework to create enhanced metabolic-regulatory network models. We used IDREAM to predict phenotypes and genetic interactions between transcription factors and genes encoding metabolic activities in the eukaryote, *Saccharomyces cerevisiae.* IDREAM models contain many fewer interactions than PROM and yet produce significantly more accurate growth predictions. IDREAM consistently outperformed PROM using any of three popular yeast metabolic models and across three experimental growth conditions. Importantly, IDREAM’s enhanced accuracy makes it possible to identify subtle synthetic growth defects. With experimental validation, these novel genetic interactions involving the pyruvate dehydrogenase complex suggested a new role for fatty acid-responsive factor Oaf1 in regulating acetyl-CoA production in glucose grown cells.

**Author Summary:** The integration of gene regulatory and metabolic network models is an important goal in computational biology, in order to develop methods that can identify the underlying mechanistic links in biological networks and advance metabolic engineering techniques. In this paper, we develop a framework called Integrated Deduced REgulation And Metabolism (IDREAM) that can improve our ability to predict phenotypes of microorganisms, and particularly it can address the challenges in evaluating phenotypic consequence of perturbing transcriptional regulation of metabolism in a eukaryotic cell. We compare the predictive performance of an IDREAM *S. cerevisiae* model with a PROM model using a TRN available from the YEASTRACT database. IDREAM outperforms PROM using any of three popular yeast metabolic models and across three experimental growth conditions, making it possible to identify subtle synthetic growth defects, and a new role for Oaf1 in the regulation of acetyl-CoA biosynthesis.

## Introduction

A major goal of systems biology is to predict the phenotypic consequences of environmental and genetic perturbations. Metabolism is a fundamental cellular system that strongly influences cell fate, and as such it is important to study the behavior and regulation of metabolic and to build models that integrate its functions with other cellular systems. Despite extensive study of the biochemistry and enzymology of metabolism for over a century, our ability to simulate the functions of metabolic networks and their interactions is still limited by their size and complexity, including their nonlinear dynamic behavior (1, 2). Traditionally, metabolic simulation has been performed using kinetic modeling, where each reaction and the dynamics of all of its components (reactants, products and enzymes) are modeled in detail. Kinetic modeling is usually applicable to small-scale biological processes and can produce accurate, dynamic predictions for fluxes, concentrations and regulatory states of the system. Kinetic modeling is limited by difficulties in parameterization, as well as the mathematical complexity of the resultant systems of differential equations. Sidestepping these size and knowledge limitations, constraint-based modeling utilizes network topology and thermodynamic constraints to make mechanistic, large-scale predictions for metabolic networks, without being dependent on detailed kinetic parameter knowledge. In recent years, the gap between kinetic and constraint-based modeling has been closing to some extent, as a number of large scale kinetic models have become available (3-6). While some of the computational limitations of kinetic modeling are being progressively overcome in well characterized systems, genome-scale models are still predominantly made using the constraint-based approach. Additionally, the dearth of publicly available, experimentally measured kinetic parameters that are necessary to populate these models, as well as their variation across different genetic polymorphisms, remains an issue. In the meantime, constraint-based modeling provides a simple, scalable, and informative method for metabolic network simulation with minimal information requirements.

Constraint-based modeling techniques (7, 8) were developed to allow researchers to simulate genome-scale metabolic networks despite these challenges, by imposing a steady state assumption. Thus, constraint-based techniques are based on computing what steady states are possible given the stoichiometry of the biochemical reaction network. Applying steady-state reaction network modeling to simulate metabolism has its roots in the 60s (9, 10), but was formalized in the 90s (11-14) under the label of flux balance analysis (FBA) (15). FBA relies on optimization techniques to identify the optimal achievable value for a particular user-defined objective in the model, such as biomass accumulation.

FBA is a powerful method for phenotype prediction due to its ability to describe stoichiometrically determined levels of substrate consumption and product production for reactions in very large metabolic systems in the absence of kinetic information or enzyme concentrations. However, one of its main drawbacks is that it does not incorporate constraints imposed upon the network by regulation of gene expression. In fact, metabolic networks are dramatically affected by complex transcriptional regulatory networks (as well as by a host of small molecule regulation processes not to be addressed herein). Changes in transcriptional regulation in response to environmental cues lead to changes in enzyme abundance or activity, which in turn lead to changes in physiological states and growth. Incorporating information about how metabolic genes are differentially regulated to metabolic network models may improve the predictions made by constraint-based analysis. The ability to integrate computational models of transcriptional regulation with models of metabolism would allow us to better describe the impact of mutations and environmental perturbations on functional metabolism. Such integrated models would have the potential to guide rational rewiring of metabolic flux and addition of new metabolic capabilities into a network (16, 17).

A common strategy for incorporating gene regulatory information into metabolic network models is to use gene expression information to impose condition-specific flux constraints on the metabolic model. This strategy depends upon the assumption that elevated gene expression measurements make it more likely that there is increased activity for the metabolic enzymes encoded by the genes with increased expression, while lower gene expression levels are more likely to correspond to lower activity of the corresponding metabolic enzymes. Methods that impose condition-specific flux constraints on metabolic network models based upon gene expression data include GIMME (18), iMAT (19), E-Flux (20), MADE (21), GX-FBA (22), MTA (23), CoreReg (24), mCADRE (25) and EXAMO (26). However, in many cases, the predictions obtained by FBA using a growth maximization objective are as good or better than those obtained using methods that incorporate gene expression to provide additional constraints (27). This discordance suggests that gene expression is not directly correlated to the activity of the encoded metabolic enzyme, or that more sophisticated methods must be employed to link gene expression data to metabolic network models. We propose that information about condition-dependent differential regulation of genes expression, such as can be captured with EGRIN (28), can provide information that can be used to improve conditional flux predictions by flux balance analysis of metabolic network models.

In previous work, some of us developed the Probabilistic Regulation of Metabolism (PROM) method for integrating transcriptional regulatory networks (TRNs) and metabolic networks (29, 30). In order to build an integrated model of a metabolic and transcriptional regulatory network for an organism using PROM, the following components are needed.

1. The genome-scale reconstruction of the metabolic network of the organism. The simulation of the metabolic network within the PROM method is performed using FBA subject to additional constraints and a penalty function.
2. A regulatory network structure, which consists of a list of transcription factors, the targets of these transcription factors, and their interactions. These transcriptional regulatory networks have generally been constructed based on high-throughput protein-DNA interaction data and/or statistical inference of functional relationships from genomic and transcriptomic data.
3. A collection of gene expression data measured under different conditions, which will allow the observation of various phenotypes for the organism under study.

PROM introduces probabilities to represent gene states and interactions between a gene and a transcription factor. In short, PROM estimates how much less an enzyme encoding gene will be transcribed when a TF is deleted and proportionally reduces the maximum flux through that enzyme. We have previously applied PROM to predict the effects of TF knockout on growth for *Escherichia coli* and *Mycobacterium tuberculosis* (29, 31). However, to date, there has not been a successful application of a PROM-like semi-automated approach to build integrative regulatory-metabolic models to predict the phenotype of TF mutants for a eukaryotic organism.

The abundance of transcriptomic data has enabled development of a number of algorithms to infer genome-scale transcriptional regulatory networks in addition to the coexpression frequency approach used in PROM (32-35). These methods have been implemented and made gene expression predictions to varying degrees of accuracy. The DREAM project (Dialogue on Reverse Engineering Assessment and Methods) evaluated over 30 network inference methods on *E. coli, Staphylococcus aureus,* and *S. cerevisiae* (36). Several methods performed relatively well for *E. coli* data sets, including CLR (32), ARACNE (37), and ANOVA (38), but not well for Yeast. Recently, several more methods were developed. RPNI (Regulation Pattern based Network Inference) defined the co-regulation pattern, indirect-regulation pattern and mixture-regulation pattern as three candidate patterns to guide the selection of candidate genes (39). Zhao et al. (40) proposed a new measure, "part mutual information" (PMI), to quantify nonlinearly direct associations in networks more accurately than traditional conditional mutal information (CMI). Another multi-level strategy named GENIMS showed better accuracy and robustness, by comparison with the methods on the DREAM4 and DREAM5 benchmark networks (40). However, significant challenges remain in accurately inferring such networks from gene expression data, particularly given the more complicated eukaryotic regulatory mechanisms in *S. cerevisiae* (36).

Environment and Gene Regulatory Influence Network (EGRIN) is an approach to meet those challenges by building a comprehensive model of condition-specific gene regulation (28). EGRIN describes which factors influence gene expression and under what environmental conditions those factors are relevant. It uses the biclustering algorithm, cMonkey (41) to find conditionally co-regulated genes from heterogeneous genome-wide datasets, and Inferelator (42) to use the mRNA expression levels of TFs or other regulators to predict the expression level of a target gene with linear regression model. EGRIN construction techniques were originally developed to study *Halobacterium salinarum* (28), but this approach was further developed for eukaryotic gene expression in the yeast *S.cerevisiae* (43). This work demonstrated that the yeast EGRIN accurately predicted condition-specific gene expression, and was able to identify transcription factors that regulate peroxisome-related genes when yeast is grown on oleic acid (43).

Here, we build upon the previous EGRIN and PROM methods to develop a framework called Integrated Deduced REgulation And Metabolism (IDREAM). IDREAM uses bootstrapping-EGRIN inferred transcriptional factor (TF) regulation of enzyme-encoding genes, then applies a PROM-like approach to apply metabolic network constraints in an effort to improve phenotype prediction, as shown in Figure 1A. We compared the predictive performance of an IDREAM *S. cerevisiae* model with a PROM model using a TRN available from the YEASTRACT database (44, 45). This comparison included growth rates predicted for TF deletion mutants, which were tested experimentally, demonstrating that predicted growth phenotypes from IDREAM were more consistent with observed phenotypes than predictions made by the PROM model. Previous work has demonstrated significant variability in growth phenotype prediction among yeast models (46), nevertheless IDREAM proved to be robust and to outperform PROM with several metabolic network models and different environmental conditions tested (Figure 2 and 4).

**Figure 1.**
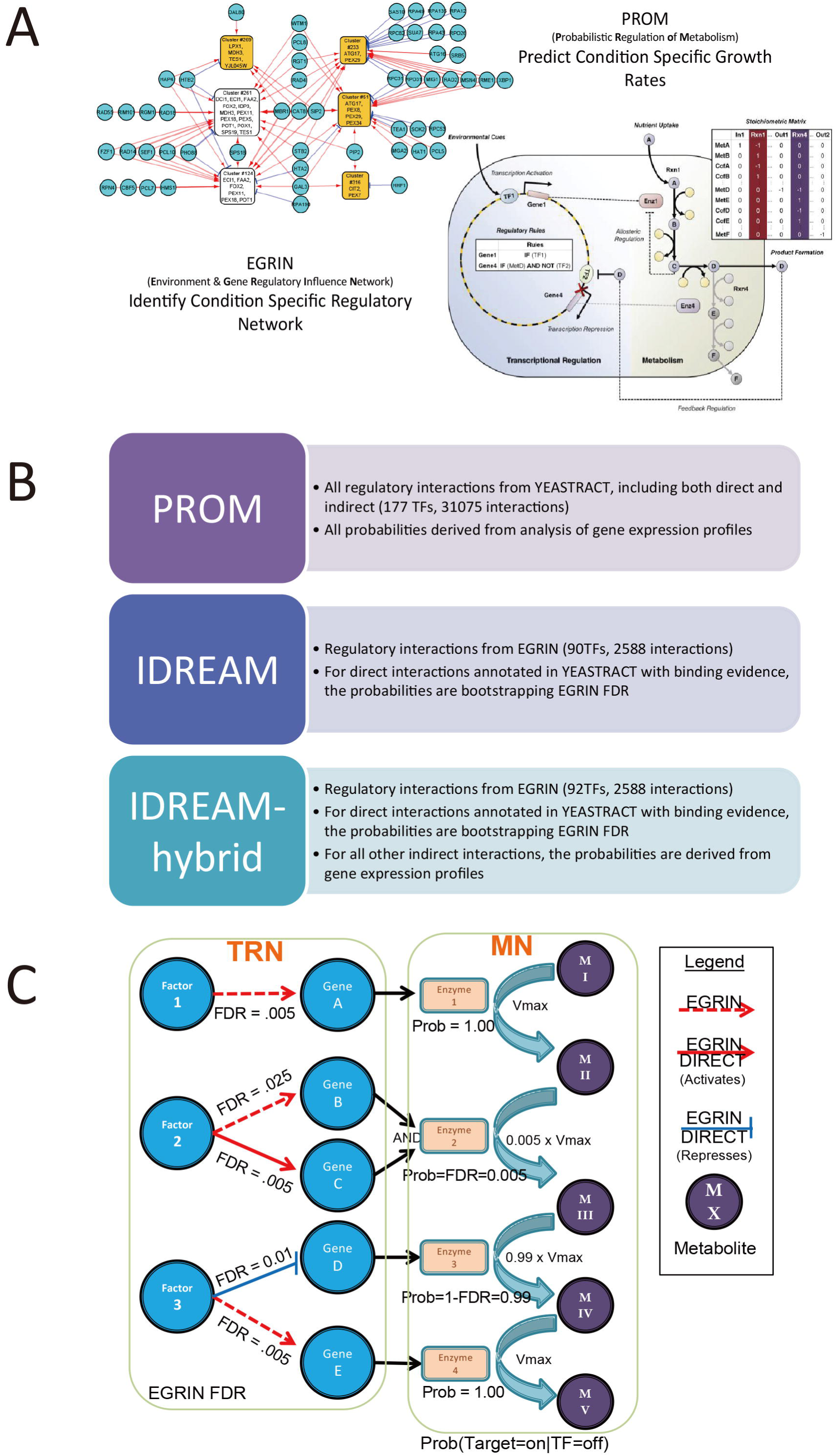
Strategy for IDREAM on integration of an EGRIN TRN with a metabolic model. A. Principle illustration of combining EGRIN and PROM for building an integrated model of a metabolic network and its corresponding gene regulatory network. B. Comparison of three integrative models: PROM, IDREAM, IDREAM-hybrid. C. Detailed illustration of probability constraints in an IDREAM model. The direct and indirect interactions are represented using solid and dashed lines, respectively. For activators (red), we set the probability to Prob(Gene=ON|Factor=OFF)=FDR. For inhibitors (blue), we set Prob(Gene=ON|Factor=OFF)=1-FDR. The constraints on the reaction flux were V_max_Prob. For indirect interactions, no effects of TF knockout on flux constraints.

**Figure 2.**
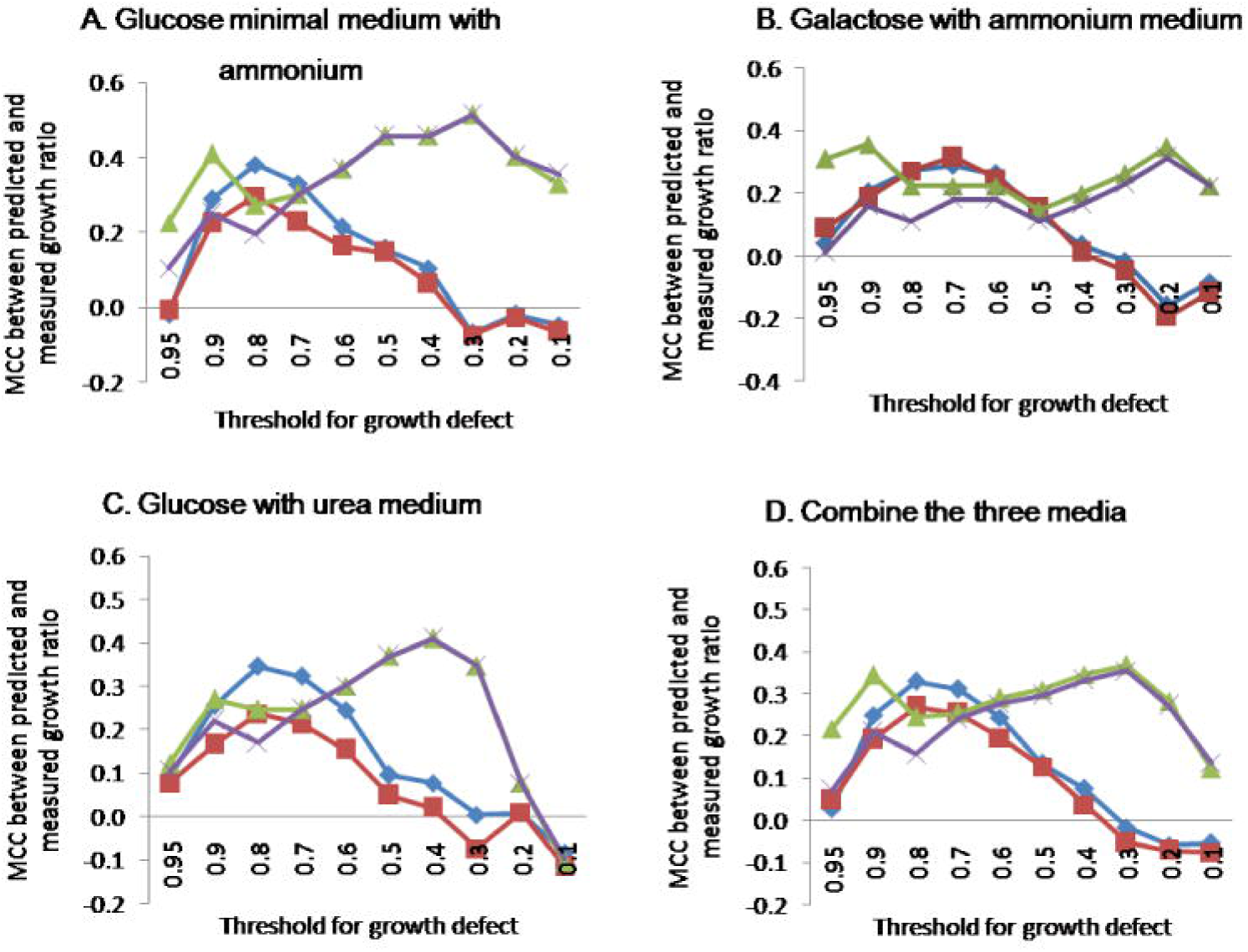
MCCs between predicted and experimental growth changes across different media and at different thresholds for binarizing a call as “growth defect” or “no growth defect.” Under each condition, we calculated the ratio of growth rates between TF knockout and wild-type. When the ratio was lower than some particular threshold, the corresponding TF is considered growth defective. By adjusting the threshold of growth ratio from 0.1 to 0.95, the MCCs between prediction and measurement were derived.

Furthermore, IDREAM enabled predictions of genetic interactions between genes encoding TFs and enzymes of the metabolic network. We experimentally tested the strongest interactions using a quantitative growth assay and validated five novel interactions between the TF Oaf1 and components of the pyruvate dehydrogenase complex. These data reveal an unexpected potential role for Oaf1 in regulating acetyl-CoA production during mitochondrial dysfunction, in addition to its well-characterized role in regulating fatty acid metabolism in the absence of glucose. Therefore, because of the inference component, the integrated network modeling approach IDREAM can uncover previously uncharacterized gene regulation of metabolism.

## Results

### Overview of the IDREAM approach for integrative regulatory-metabolic modeling

The original PROM framework represents the TF influence with a conditional probability derived from analysis of gene expression profiles. This conditional probability estimates the likelihood that an ON/OFF state in a TF will lead to an ON/OFF state in the target genes (29). For IDREAM, the conditional probability was instead represented by the bootstrapping EGRIN-derived FDR values for the subset of EGRIN-discovered regulator interactions that also have evidence for direct interaction in the YEASTRACT database. As shown in Figure 1B and 1C (see details in Methods), this allows us to represent the TF influence, while leaving the rest of the metabolic reactions unconstrained by regulation. Essentially, IDREAM focuses on our highest confidence set of interactions, where there is both evidence for direct regulation from YEASTRACT and a strong transcriptional influence that is sufficient to be predicted by the inference techniques of EGRIN. In addition, we generated an IDREAM-PROM hybrid model, which is able to adjust the conditional probabilities for the indirect interactions, using the conventional PROM approach coupled with the IDREAM constraints on the high confidence. As above, EGRIN inferred interactions also have evidence for direct interaction in the YEASTRACT database. We compared these two integrated models with a standard PROM model that solely uses interactions from the YEASTRACT database as the regulatory component, without any information from the EGRIN regulatory network model. The strategies for construction of the three integrative models are described in Figure 1B. For all the approaches, we used Yeast6 as the base metabolic model (47). Yeast6 was found to be as accurate as any available yeast reconstruction for growth predictions based on an extensive metabolic comparison across all published models and available datasets (46), and also performs best with IDREAM. We compared the network properties and the predictive performance of the resulting three integrative models: PROM, IDREAM, and IDREAM-hybrid. To test the effectiveness of these integrated models, we validated the growth predictions against growth rate data for 119 TF knockouts measured by the Sauer Laboratory (48).

The standard PROM model included 177 TFs and a total of 31,075 regulatory associations from YEASTRACT, of which 7,292 were direct interactions with evidence of TF binding. By mapping the target genes in the TRN with metabolic genes in the MN, we integrated 2588 EGRIN-inferred influences consisting of 91 TFs transcriptionally regulating 794 genes encoding enzymes of the metabolic network with false discovery rates (FDR) ≤ 0.05 (See Methods for details). There were 307 interactions in the IDREAM model annotated as direct regulatory associations in YEASTRACT for which evidence of TF-chromatin binding has been generated. Although there are many more TFs and interactions in YEASTRACT, 15 out of the 17 TFs observed to cause growth defects upon deletion (48) are included in the EGRIN network (shown in Figure S1). Additionally, for the total 900 genes encoding enzymes of the metabolic network in the Yeast6 model, PROM and IDREAM included 863 and 794 genes respectively, which suggested that the regulatory network generated by EGRIN captures the part of the network that is most relevant to phenotypic predictions influenced by changes in metabolic flux, while sparing extraneous components.

### IDREAM predicted growth phenotypes with significantly better accuracy than PROM

Predicting gene essentiality is a basic and important task for genome-scale metabolic models (49-51). Advanced models that include TF regulators of genes encoding metabolic enzymes (such as IDREAM and PROM) can also predict growth rates when TFs are deleted. We used FBA to calculate the optimal growth rate on glucose-containing minimal medium using the Yeast6 model. Then, using the three regulatory-metabolic models, we simulated the growth rate for each TF knockout. The ratio of mutant vs. wild-type growth rate was compared with the growth ratio for 119 TF knockouts previously measured (48). There are 90 TFs and 52 TFs with corresponding deletion mutant growth ratios in the PROM and IDREAM models, respectively. There were 51 TFs in common between the two integrative models, so we distinguish PROM by TF90 (the whole YEASTRACT-based model) and TF51 (the portion of the YEASTRACT-based model that overlaps with that from IDREAM). As shown in Table 1, the Pearson Correlation Coefficient (PCC) between experimental results and predictions by IDREAM is much higher than that by PROM (PCC is 0.43 vs. 0.17), and the normalized sum of squared error is significantly lower for IDREAM (0.12 vs. 0.25). We performed a two-tailed t-test testing the null hypothesis that the mean absolute residuals for IDREAM are the same as the mean absolute residuals for PROM, and obtained p-value=0.01 (Table 2). Interestingly, the performances of IDREAM and IDREAM-hybrid were very similar, suggesting that the core set of direct regulatory interactions predicted from the EGRIN approach plays a key role in affecting phenotype, irrespective of the conditional probabilities calculated for the indirect interactions.

**Table 1.** Comparison of PROM and IDREAM predicted growth ratio with experiments under glucose minimal medium. The ratio of mutant vs. wild-type growth rate was compared with the growth ratio for 119 TF knockouts previously measured by Sauer Lab. There were 51 TFs in common between the two integrative models, so we distinguish PROM by TF90 (the whole YEASTRACT-based model) and TF51 (the portion of the YEASTRACT-based model that overlaps with that from IDREAM).

| Integrative model | Correlation | p-value | Sum of squared error | Normalized sum of squared error / permutation p-value |
| --- | --- | --- | --- | --- |
| PROM_TF90 | 0.2110 | 0.0459 | 4.298 | 0.205 / 0.029 |
| PROM_TF51 | 0.1712 | 0.2297 | 3.566 | 0.249 / 0.144 |
| IDREAM-hybrid | 0.4183 | 0.0020 | 2.481 | 0.118 / 0.004 |
| IDREAM | 0.4325 | 0.0014 | 2.506 | 0.121 / 0.003 |

**Table 2.**
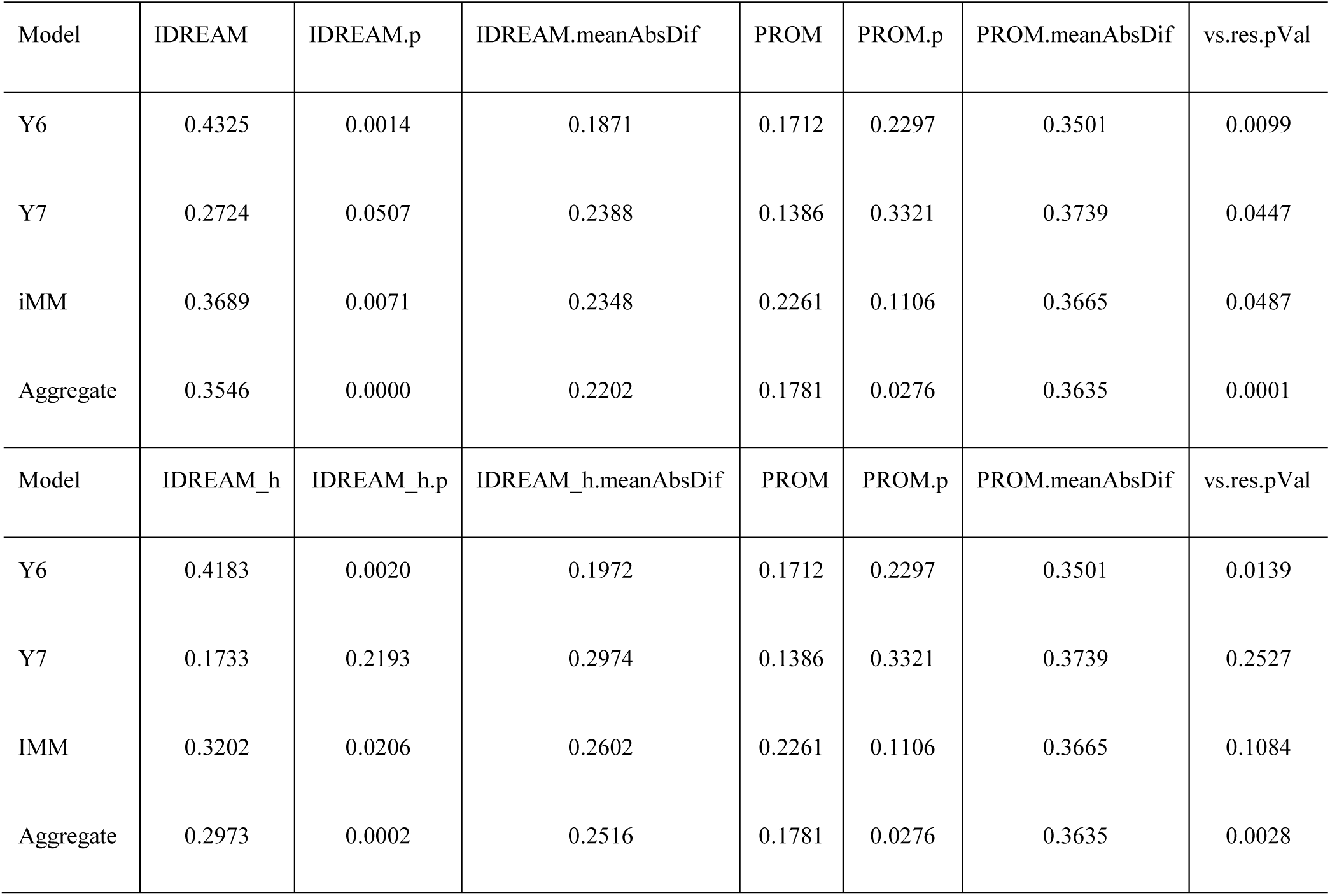
Comparison of mean absolute residuals for IDREAM and PROM aggregating different yeast models. The first column shows three different yeast metabolic models, aggregate refers to the predictions for all three models taken together. Column 2-4 show the Pearson correlation coefficient, p-value, and mean absolute residuals difference between predicted and actual growth by IDREAM and IDREAM_hybrid model. Column 5-7 show the Pearson correlation coefficient, p-value, and mean absolute residuals difference by PROM_TF51. Column 8 ‘vs.res.pVal’ represents the significance of difference in correlations between the two IDREAM models and the PROM model. P-values were calculated using a Fisher’s Z transform. IDREAM_h means the IDREAM_hybrid model.

| Model | IDREAM | IDREAM.p | IDREAM.meanAbsDif | PROM | PROM.p | PROM.meanAbsDif | vs.res.pVal |
| --- | --- | --- | --- | --- | --- | --- | --- |
| Y6 | 0.4325 | 0.0014 | 0.1871 | 0.1712 | 0.2297 | 0.3501 | 0.0099 |
| Y7 | 0.2724 | 0.0507 | 0.2388 | 0.1386 | 0.3321 | 0.3739 | 0.0447 |
| iMM | 0.3689 | 0.0071 | 0.2348 | 0.2261 | 0.1106 | 0.3665 | 0.0487 |
| Aggregate | 0.3546 | 0.0000 | 0.2202 | 0.1781 | 0.0276 | 0.3635 | 0.0001 |
| Model | IDREAM_h | IDREAM_h.p | IDREAM_h.meanAbsDif | PROM | PROM.p | PROM.meanAbsDif | vs.res.pVal |
| Y6 | 0.4183 | 0.0020 | 0.1972 | 0.1712 | 0.2297 | 0.3501 | 0.0139 |
| Y7 | 0.1733 | 0.2193 | 0.2974 | 0.1386 | 0.3321 | 0.3739 | 0.2527 |
| IMM | 0.3202 | 0.0206 | 0.2602 | 0.2261 | 0.1106 | 0.3665 | 0.1084 |
| Aggregate | 0.2973 | 0.0002 | 0.2516 | 0.1781 | 0.0276 | 0.3635 | 0.0028 |

In order to more fully evaluate the generality of these results, we expanded our set of predictions for two additional growth conditions measured previously (48): galactose with ammonium as a nitrogen source, and glucose with urea as a nitrogen source. Additionally, across all three growth conditions, we evaluated the effect of changing the cutoff for binarizing the data into categories of “growth defect” vs. “no growth defect”. We utilized the Matthews correlation coefficient (MCC) (52) as it is the common method of choice for statistically assessing performance of binary classifications. The MCC results for the two IDREAM predictions were much better than those for PROM overall (Figure 2). In particular, when the threshold of ratio for growth defect was less than 0.5, the Fisher’s transformation test for the pairs of MCC values showed that IDREAM significantly outperformed PROM across all measured conditions (p <0.05). IDREAM can also decrease the variability of the estimated internal fluxes in addition to predicting reaction essentiality for growth. Applying flux variability analysis (FVA) to the IDREAM model can reveal predicted changes in the reaction flux solution space that result from each TF perturbation. Comparison with the FVA results of the original YEAST6 model showed that the solution space is reduced for IDREAM (Table S1).

We also estimated the significance of the predictive performance by randomly permuting the expression data and TF-gene associations. For the expression dataset, we fixed the number of genes and randomly permuted the expression values 500 times, and then calculated the percentage of permutations that generated higher MCCs than the constructed IDREAM model (designated as a p-value). Additionally, we generated 500 permutated regulatory networks by fixing the number of TFs and genes and randomly permuting their connections, while the expression dataset remained unchanged. The percentage of permuted networks that generated higher MCCs than the constructed IDREAM model was calculated as a p-value. We found that the MCCs from the IDREAM and IDREAM-hybrid model were all significant against the distribution of permutations of expression and network associations (p < 0.05 in each case) (Table S2). The predictive accuracy of the EGRIN-derived TF regulatory influences on metabolism was further underscored by the observation that an integrated model that was constructed by integrating TF influences inferred by CLR (36) made growth rate predictions that did not correlate with experimental data (Table S3).

To further evaluate the performance of IDREAM compared to PROM, and because growth ratios are continuous values, we tested whether PROM or IDREAM performed better at predicting a range of growth defects. Thus, instead of considering only 50% ratio as defining a growth defect, we considered multiple threshold ranging from severe growth defects (~10% of WT) to virtually normal growth rates and compared the performance of IDREAM and PROM using Receiver Operator Characteristic curves (53) (Figure 3, Figure S2 and Table S4). Overall, the mean Area Under the Curve value (54) for this wide range of thresholds was significantly higher for IDREAM than PROM (0.67 vs 0.58, Wilcoxon signed rank p-value < 0.004; Figure S2, Table S4) indicating that IDREAM more accurately predicted growth defects.

**Figure 3.**
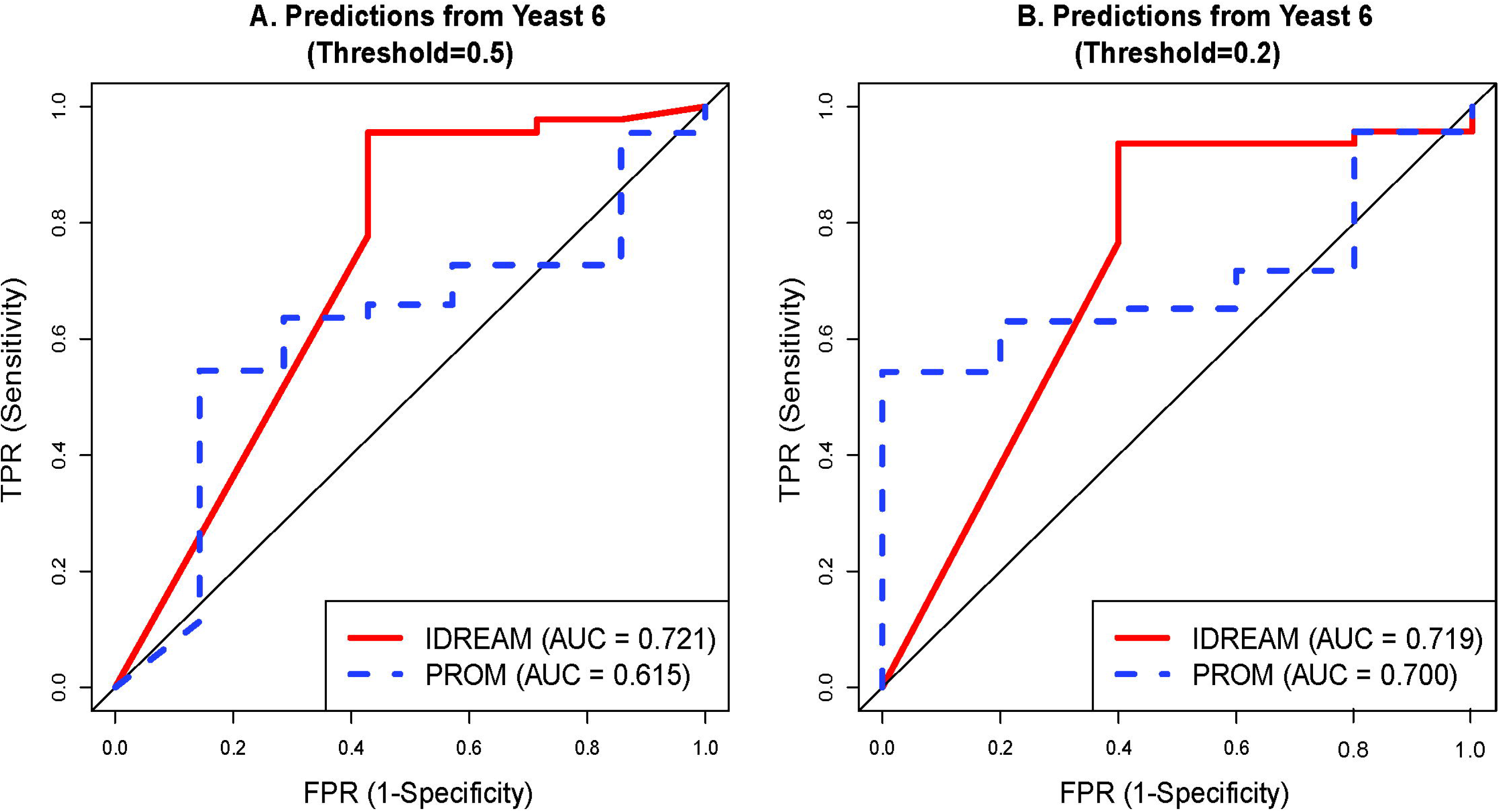
ROC curves for growth defect predictions using IDREAM and PROM on Yeast6 model. A. Threshold is 0.5 for binarizing a call as “growth defect” or “no growth defect” B. Threshold is 0.2 for binarizing a call as “growth defect” or “no growth defect”

### IDREAM outperforms PROM for different yeast metabolic models

To further validate the performance of IDREAM, we tested our approach with two metabolic reconstructions of yeast other than Yeast6: Yeast7, the latest published reconstruction of yeast (55), and iMM904 by the Palsson Lab (56), probably the most widely used reconstruction. We compared the MCC between predicted and experimental growth ratios for three representative thresholds (0.2, 0.5, and 0.95) by binarizing a call as either ‘growth defect’ or ‘no growth defect’. As shown in Figure 4, the MCCs for IDREAM were larger than those for PROM for all three metabolic models, especially for thresholds of 0.2 and 0.95. Although the MCCs for Yeast6 were larger for most thresholds, there was no significant difference for ROC curves among the three reconstructions (Figure S3A). Also, the ROC curves produced by PROM for the three metabolic models did not show significant differences (Figure S3B), but the AUC values for IDREAM were generally higher than those for PROM. The PCC between predicted and experimental growth ratio by the three models also demonstrated that IDREAM outperformed PROM, two-tailed p-values testing the mean absolute residuals were (in aggregate) significant (p-value < 0.05) across all metabolic models, as shown in Table 2.

**Figure 4.**
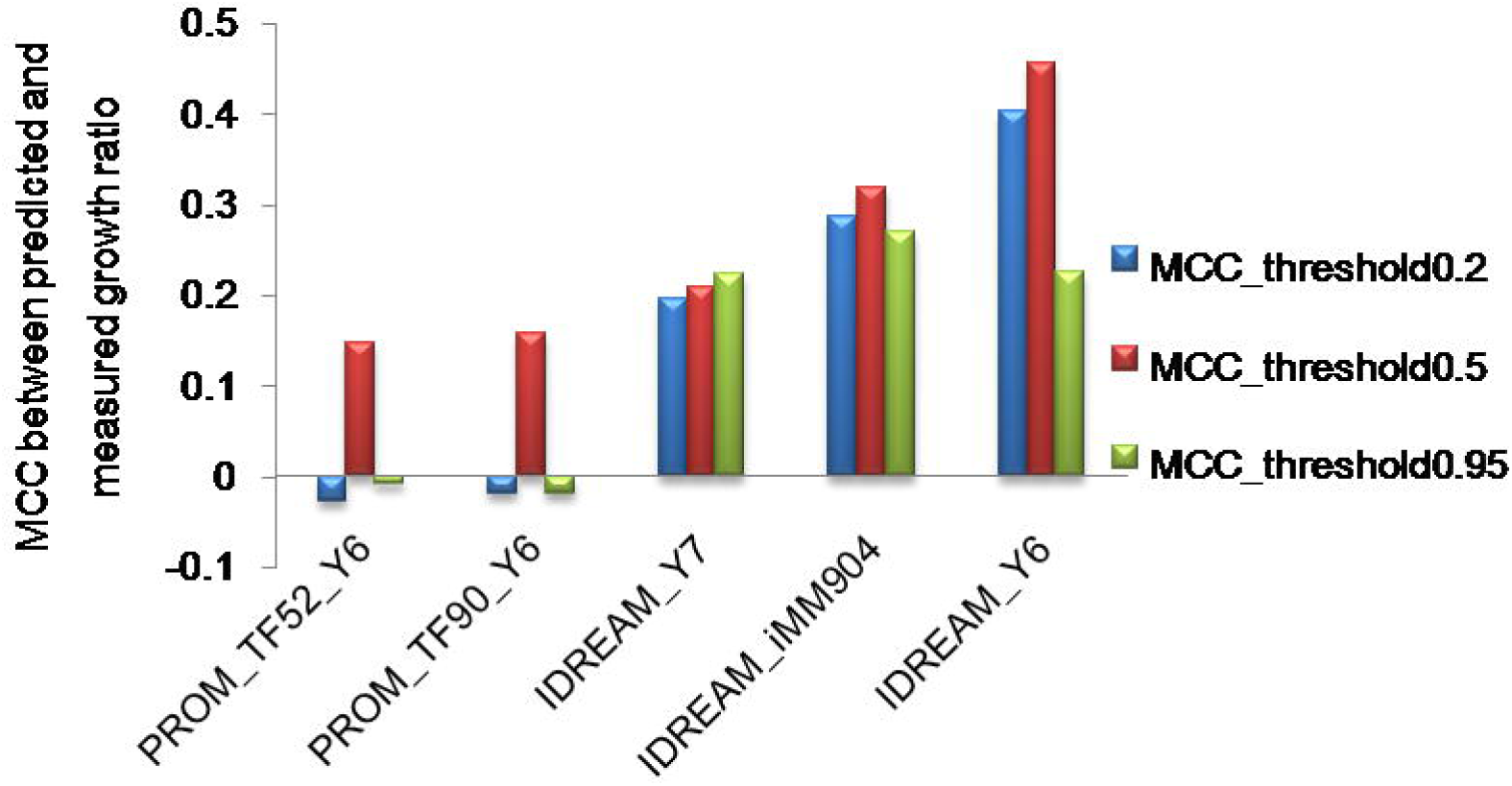
MCCs by different integrative models using different thresholds of growth ratio determining growth defect. Y6, Y7, and iMM904 refer to the Yeast metabolic models Yeast 6, Yeast 7, and iMM904 respectively.

We also predicted the growth ratios for the three reconstructions across different conditions, calculated the Pearson's correlation to experimentally determined growth ratios, and determined p-values based on the Fisher’s Z transform (see Methods). As shown in Table S5, the aggregate correlation for IDREAM predictions was significantly higher than that for PROM or for IDREAM-hybrid. We conclude that phenotypic predictions were significantly better with IDREAM, whether analyzed with Matthews or Pearson correlation.

### IDREAM model effectively predicted phenotypes of double gene deletions

Using the IDREAM model, we further simulated the growth phenotypes of strains with double-deletions of genes encoding a TF paired with a gene encoding an enzyme of the metabolic network, as shown in supplemental Figure S4. The model predicted a dramatically reduced growth rate for several double deletion strains, but predicted no growth defects for the corresponding single deletion strains. Thirty-nine such pairs were predicted to vary by over 90% when comparing predictions for single and double deletion growth rates (see detail in Methods). Figure 5 shows the predicted interacting pairs with the most dramatic reduction in predicted growth rates for the double deletion mutants (> 95% less than each single deletion). For these, deletion of either the TF or metabolic gene individually had no predicted effect on growth, but the double deletion resulted in a predicted growth rate of zero. These predictions are based on global mRNA levels in response to gene deletions and other perturbations, with the assumption that mRNA levels reflect abundance or activities of their encoded proteins. If mRNA levels were perfectly matched with protein levels, we would consider such gene pairs as predicted synthetic essential (57); but as mRNA and protein levels are only partially correlated (58), we instead consider these pairs to be candidate negative/aggravating interacting pairs.

**Figure 5.**
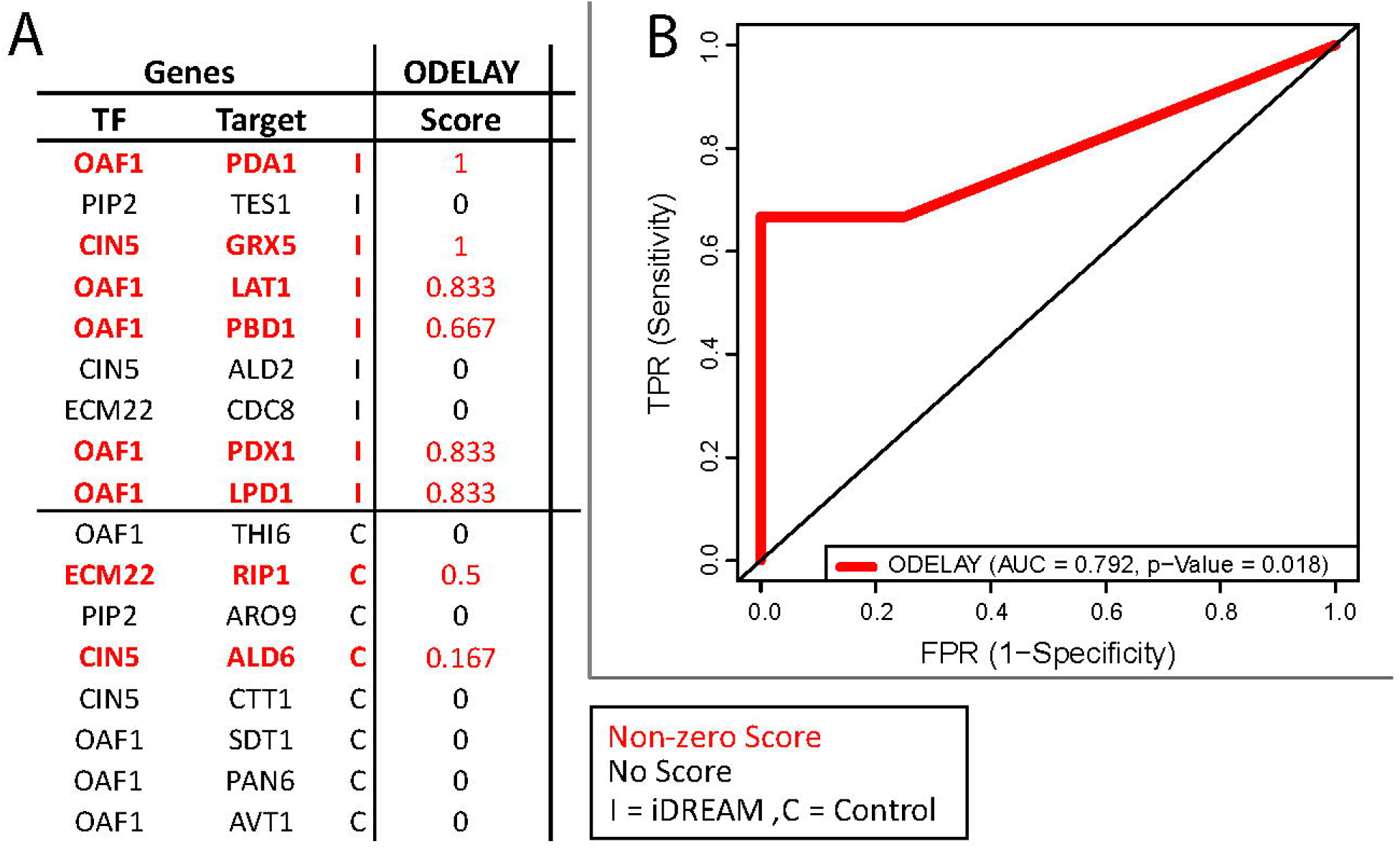
Synthetic growth defect interactions identified by IDREAM. A. Growth defect confidence scores measured by ODELAY. Beyond the genetic interactions between OAF1 and genes encoding components of the pyruvate dehydrogenase (PDH) complex, ODELAY also validated the predicted genetic interaction between CIN5 and GRX5, but did not confirm the remaining 3 of the 9 predictions. B. ROC curve describing identification of IDREAM or control strains based on ODELAY scores.

These 9 predicted genetic interactions were tested experimentally along with 8 control pairs (consisting of the same TFs and randomly selected genes encoding metabolic enzymes where the double deletion was predicted to have no synthetic defect) (Figure 5, Figure S5). We used a quantitative technology called ODELAY (One-cell doubling evaluation by living arrays of yeast) (59) to test each double deletion strain in the presence of 2% glucose, which can track the growth of many individual colonies for each strain over time using high resolution imaging (Figure 5A). The method yields a measure of doubling times for each clone in a population. Synthetic interactions are revealed when the growth defect of the double mutant is greater than the sum of each single mutant. The quantitative approaches showed genetic interactions between OAF1 and five genes encoding components of the pyruvate dehydrogenase (PDH) complex including LAT1, PDA1, PDB1, PDX1 and LPD1. Beyond these genetic interactions between OAF1 and genes encoding components of the pyruvate dehydrogenase (PDH) complex, ODELAY also validated the predicted genetic interaction between CIN5 and GRX5 but did not confirm the remaining 3 of the 9 predictions. Overall, it demonstrated that IDREAM made accurate predictions of synthetic interactions among gene pairs (Figure 5B, AUC = 0.792, Mann-Whitney p-value = 0.018).

## Discussion

In this study, we developed and applied an approach, IDREAM, which integrated together a network inference algorithm (EGRIN) into the previous constraint-based regulatory-metabolic modeling framework (PROM) to build a combined gene regulatory-metabolic network model for yeast. The major outcomes of this study were (1) the prototyping of the IDREAM approach; (2) demonstrating superior performance of IDREAM compared with PROM across a variety of metrics, where the latter approach had not been successful in building a combined gene regulatory-metabolic network model for a eukaryotic cell; (3) demonstrating direct interaction sets and activation/inhibition status are important factors for generating accurate predictions with IDREAM; (4) predicting genetic interactions, including across joint TF and enzyme perturbations. Importantly, these predictions were experimentally validated, both using existing gene knockout essentiality and growth rate information, as well as in a set of experimental results generated herein and in quantitative growth assessments in the yeast mutants using ODELAY (59). Each of these points will be discussed in detail in the following.

### Integration of an inferred regulatory network with a constraint-based metabolic model

Integration of a gene regulatory network with a metabolic network at genome-scale poses significant challenges, in part because they are distinct network types requiring very different modeling frameworks. While the PROM framework integrates regulatory and metabolic networks at genome-scale, the type of regulatory interactions it has incorporated have typically been limited to those that are supported by physical evidence such as from ChIP-chip/Seq experiments (29). Since a comprehensive map for protein-DNA (P-D) interactions of all TFs and their targets is not typically available for most organisms, this greatly limits the general utility of PROM. Even when they exist, the P-D interaction map for any given organism is incomplete as the interactions are typically mapped in one or few environmental conditions, and all interactions may not have causal consequences on metabolism. EGRIN overcomes this limitation by discovering direct and indirect causal regulatory influences of TFs that act in an environmental condition-dependent manner on their downstream target genes (28, 43). With IDREAM, we have demonstrated an approach that integrates regulatory influences learned from EGRIN to augment the previous PROM approach for building integrated metabolic-regulatory network models. This approach led to accurate predictions of growth-altering synthetic interactions across the regulatory and metabolic network of *S. cerevisiae.* IDREAM is generalizable to any organism with a sequenced genome, reconstructed metabolic model, and sufficient gene expression data. Accuracy of phenotype predictions by IDREAM were significantly better with EGRIN relative to when regulatory interactions from the CLR method was integrated (32). This result demonstrated the importance of incorporating indirect causal influences in accurate phenotype prediction by IDREAM, as the CLR method considers only previously known and mostly direct regulatory interactions supported by evidence of physical P-D interaction of the TF and its target gene promoter (Table 1 and Table S3).

### The integrative IDREAM model predicted phenotypes better than the PROM model

One of the important roles of constraint-based models is to predict which genes encoding metabolic enzymes are essential for growth in a particular environmental condition, given a set of nutritional inputs. Here, we expanded the scope of our model through integrative regulatory-metabolic modeling of the effects of TF knockout on growth. The Pearson correlation coefficient between predicted and experimental growth ratios for IDREAM was significantly higher than that for PROM (Table 2: PCC=0.43 vs. 0.17 respectively, p-value=0.01). However, since there was no linear relationship in the distribution of growth ratio for each TF mutant, we also computed the Matthews Correlation Coefficient for model predictions by setting different growth ratio thresholds for categorizing gene deletion strains that have ‘growth defect’ or ‘no growth defect’. Overall, MCCs were much larger for IDREAM-predictions relative to PROM, especially when the growth ratio cutoff was less than or equal to 0.5 (Figure 2). IDREAM also outperformed the standalone metabolic model at predicting gene essentiality (Figure 3): the standalone metabolic model cannot predict the effects of TF knockouts, therefore we assume its ROC curve is the same as random (the diagonal line in Figure 3).

To examine whether these predictions were sensitive to a particular metabolic reconstruction, we tested IDREAM performance with three distinct models: the consensus reconstructions Yeast6 (47), Yeast7 (55), and iMM904 (56). Although Yeast6 generated better correlations across several growth ratio thresholds (Figure 4), the AUC for the ROC curves was similar across the three metabolic models. Importantly, IDREAM performed better than PROM, regardless of which metabolic model was used (Figure S3). These comparisons demonstrate that IDREAM is significantly better at uncovering the influence of regulation on downstream phenotypes, irrespective of the version of the reconstructed metabolic model.

### Direct interaction sets and activation/inhibition status are important factors for generating accurate predictions

The original PROM method used a gene regulatory network structure from public resources (such as YEASTRACT), including both direct and indirect interactions, and the probabilistic influence for these two different interaction sets was calculated from gene expression correlations between the TFs and their target genes. However, there was poor correlation between PROM model predictions and observed growth ratios (PCC=0.17, p-value=0.23). The correlation was even worse when we restricted the PROM model to include just the 7,292 interactions with binding evidence in YEASTRACT (PCC=0.076, p-value=0.48). In contrast, there was significant correlation between observed phenotypes and IDREAM model predictions when constraints derived from EGRIN were applied to a core set of regulatory interactions with binding evidence (direct interactions) from YEASTRACT. This correlation was significant whether the TF influences of indirect interactions were constrained using PROM as is done for IDREAM-hybrid (PCC=0.42, p-value=0.002) or left unconstrained (PCC=0.43, p-value=0.001). Thus, constraints on direct interactions using EGRIN-derived FDR produces much better TF knockout phenotype predictions by IDREAM relative to the standalone PROM approach. Growth rate prediction by IDREAM was further improved by accounting for the activator and inhibitor status of TFs and by using the bootstrapped-EGRIN FDR to guide the probabilistic influence of TFs on their target metabolic genes. These results demonstrated that PROM may overlay unnecessary constraints on indirect TF interactions, and may therefore erroneously predict that a TF deletion will result in decreased growth rate. In contrast, IDREAM can differentiate direct and indirect interactions, and furthermore identify the high confidence TF-interactions that have both evidence of direct regulation from YEASTRACT and EGRIN-predicted influence on downstream target genes.

### Predicted genetic interactions with OAF1 were validated and relevant to acetyl-CoA regulation

The IDREAM-predicted negative interactions of OAF1 (encoding TF Oaf1) with genes PDX1, PDA1, PDB1, LPD1, and LAT1 in the presence of glucose were validated experimentally (Figure 5). These 5 genes encode components of the PDH complex, a mitochondrial enzyme that generates acetyl-CoA from pyruvate. Acetyl-CoA has important roles in various aspects of cell biology and its metabolism is compartmentalized and tightly regulated.

These predictions were initially surprising as Oaf1 functions primarily in presence of fatty acids to up-regulate genes involved in peroxisome biogenesis and function, including β-oxidation (60). By contrast, glucose represses Oaf1-mediated activation (61). However, acetyl Co-A can also be produced in peroxisomes (62), from where it is exported to the cytoplasm for a diverse array of functions in the TCA cycle, amino acid and carbohydrate biosynthesis (63), and cell signaling (64). It is likely that Oaf1 plays a role in regulating acetyl CoA production in peroxisomes and compensates for a dysfunctional PDH complex during growth in the presence of glucose. This regulatory interaction is further supported by flux balance analysis, which predicted that biomass production is lower when both the PDH complex and acetyl-CoA producing reactions in the peroxisome are active, relative to biomass produced when only one of these pathways is active. In sum, these data suggest that Oaf1-mediated control of alternate pathways for acetyl-CoA production has significant influence on biomass production.

This hypothesis implicates communication between the mitochondrion, where the PDH complex is localized, peroxisomal acetyl-CoA production and the nucleus, where Oaf1 controls transcription. Such communication is evident by systems level studies demonstrating coordinated activities between peroxisomes and mitochondria (61), shared and differential localization of peroxisomal and mitochondrial proteins (63), and the discovery of retrograde signaling molecules controlling communication between peroxisomes, mitochondria and the nucleus (65, 66). Thus, in response to mitochondrial PDH complex dysfunction, peroxisomes could export acetyl-CoA via the carnitine shuttle or export glyoxylate pathway intermediates such as citrate to the cytoplasm (62). Indeed, peroxisomal citrate synthase (CIT2) is up-regulated in the retrograde response (67), and CIT2 has negative genetic interactions with components of the PDH complex (PDA1, PDB1, PDX1, and LAT1) (68).

Additionally, YAT2, one of three carnitine acetyltransferases in *S. cerevisiae,* is dramatically upregulated in an OAF1 deletion in the presence of glucose (5.2 fold, p-value 0.01773) (34).

OAF1 deletion and PDH dysfunction form synthetic lethal pairs which can be understood by analyzing the glyoxylate pathway. Deletion of OAF1 results in moderate downregulation of 4 of 5 glyoxylate metabolic genes including MDH3, CIT2, ACO1, and ICL1 during growth in glucose (34), suggesting Oaf1 normally has a role in promoting the expression of these genes. This suggests that in the absence of Oaf1, cells may be poorly suited to increasing the export of glyoxylate pathway intermediates due to reduced expression of glyoxylate metabolic genes. Oaf1 could also function by upregulating PEX genes involved in peroxisome biogenesis, which could affect localization of peroxisomal metabolic enzymes. Data show that deletion of OAF1 results in reduced expression of 21 of 27 PEX genes measured including greater than 2-fold downregulation of PEX3, PEX12, PEX13, PEX17, PEX19, and PEX34 during growth in 2% glucose (34). Consistent with this, 13 negative genetic interactions have been found between PEX genes and components of PDH complex (68) and peroxisomes have been shown to proliferate under conditions of mitochondrial dysfunction (69).

## Conclusion

In conclusion, the IDREAM approach demonstrates that it is possible to predict phenotypic consequence of perturbing transcriptional regulation of metabolism in a eukaryotic cell. This predictive capability of IDREAM revealed a new role for Oaf1 in the regulation of acetyl-CoA biosynthesis, exposing the phenotypic consequence of combinatorial perturbations to this regulatory-metabolic network during growth on glucose. It is notable that IDREAM is capable of making reasonably accurate predictions without explicitly modeling many additional layers of control, such as allosteric regulation and post-translational protein modification, that are known to establish important mechanistic linkages between transcription and metabolism (17). While this capability of IDREAM to predict flow of information from transcription->metabolism->phenotype is powerful and useful for directing laboratory experiments, it is important to integrate and model the intervening regulatory processes in order to identify the mechanistic linkages and advance metabolic engineering.

## Materials and Methods

### Yeast regulatory network inferred using EGRIN

We expanded the yeast gene regulatory network derived using EGRIN and presented in (43), in order to integrate it with PROM by focusing on predicting regulation for individual genes rather than for gene clusters as had been done previously. The yeast EGRIN was constructed using two computational tools (cMonkey and Inferelator) trained considering 5939 yeast genes in 2929 microarray experiments and evaluating 392 of those genes as possible regulators (i.e. factors). cMonkey identified biclusters of genes that were coherently expressed in some of these experiments, while Inferelator identified regulators of those genes by using (hybrid) linear models (42). To improve the gene level predictions over those in the previously published yeast EGRIN, we made the Inferelator regression more robust by generating additional linear models. For each of the 5939 target genes, we constructed separate models from 200 randomly selected subsets of the 2929 experiments, as well as a 201^st^ model constructed using the entire data set. This resulted in 201 generated gene regulatory models for each of the 5939 yeast genes, for a total of 1,193,739 models. For each gene, we estimated a false discovery rate (FDR) for each factor by tallying the fraction of models that identified that factor as a regulator. Thus, if factor X was predicted to regulate gene Y in 191 of 200 models, then X would have an FDR=1–191/200 = 0.045. We included only those interactions that passed an FDR cutoff of 0.05 and interpreted the remaining FDRs such that the fraction of times that a factor was predicted to regulate a target corresponded to the fraction of that targets activity that was not controlled by that regulator. Therefore, if X is predicted to activate Y with an FDR of 0.045, only 4.5% of Y’s activity would be predicted to remain if X was deleted. If X is predicted to deactivate Y, then we use the much larger 1 - FDR (e.g., 95.5% of activity) to represent that Y is somehow disturbed without a significant reduction in activity. We predicted whether a factor was an activator or repressor by testing if its mRNA expression was correlated or anti-correlated (respectively) with the expression of its target under the relevant experimental condition. The interactions between TFs and target genes in EGRIN and YEASTRACT TRN are listed in Table S6.

### Yeast metabolic model and flux balance analysis

The genome-scale metabolic model for yeast has been updated through iterative collaborative curation by multiple research groups. We downloaded the yeast consensus reconstruction (70) versions 6.06 (47) and 7.01 (55) from the SourceForge repository (http://yeast.sf.net/), and acquired the iMM904 model from (56).

We used the COBRA Toolbox (51) to conduct FBA. Briefly, FBA is a mathematical optimization method for calculating a maximum or minimal achievable metabolic flux, subject to the constraints imposed by metabolic network stoichiometry, thermodynamic information, and capacity constraints (51).

### IDREAM integrative model construction

We constructed the IDREAM model by using the inferred EGRIN regulatory network to constrain reactions in yeast metabolic network models. There were 307 interactions in the EGRIN network annotated as direct regulatory associations in YEASTRACT that have binding evidence. If the deleted TF is an activator with binding evidence, the probability of a target gene being ON was set as the bootstrapping Inferelator-derived FDR, i.e. Prob(Gene=ON|Factor=OFF)=FDR. If the deleted TF is an inhibitor with binding evidence, we set Prob(Gene=ON|Factor=OFF)=1-FDR, as shown in Figure 1B. In contrast, the TF influences of indirect interactions were unconstrained (IDREAM) or had conditional probabilities inferred using the expression datasets (IDREAM-hybrid). Then the constraints on the corresponding reaction flux were V_max_ Prob, where V_max_ was derived by flux variability analysis (and thus represents the effective Vmax based on constraints throughout the network). The implementation of the IDREAM method for yeast can be downloaded as supplemental Script S1.

### Experimental growth rate for TF knockouts in *Saccharomyces cerevisiae*

Fendt et al. (48) systematically measured growth rates in 119 transcription factor deletion mutants of *Saccharomyces cerevisiae* under five growth conditions. Since low pH and high osmolarity cannot be simulated with FBA, we took the growth rates of 119 mutants under three conditions: glucose with ammonium as nitrogen source, galactose with ammonium as nitrogen source, and glucose with urea as nitrogen source.

### Matthews correlation coefficient for evaluation of gene essentiality prediction

The agreement between model gene essentiality predictions and the reference lists was quantified using the Matthews Correlation Coefficient (equation 1) (52), a metric that considers true positive, true negative, false positive, and false negative predictions without any assumption of the frequency of observations in the reference dataset. MCC ranges from −1 (when model predictions are the exact opposite of the reference dataset) to +1 (when model predictions match the reference data set).

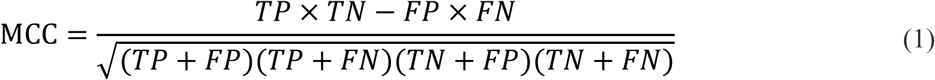

Where true positives (TP), true negatives (TN), false positives (FP), and false negatives (FN) are defined based on the measurement by (48). A true positive prediction is one in which the model predicts that a gene is essential for growth, and the gene has also been annotated as essential.

### Pearson correlation coefficient for evaluation of growth predictions

Experimentally determined yeast growth rates that have been normalized to wild-type growth rates (48) were compared to growth rates predicted by the metabolic models. The reported Pearson product-moment correlation (71) measures the linear correlation between the predicted and experimentally determined growth rates for yeast strains. Aggregate predictions were made by concatenating the lists of predictions for all three models and comparing to the appropriate experimentally determined growth rates. Thus, if each of the three models makes a different growth rate prediction for the same yeast strain, then that yeast strain will be represented three times in the aggregate calculation.

### Prediction of interacting pairs of genes encoding TFs and metabolic genes

Synthetic lethality or sickness occurs when the combination of two gene deletion results in reduced fitness and can identify buffering relationships where one gene can compensate for the loss of another (57). We predicted these synthetic relationships based on the variation of growth rates between single and double deletions. We calculated the difference in growth rates between single_TF_deletion and double_TF_gene_deletion, represented as Diff1, and the difference in growth rates between single_gene_deletion and double_TF_gene_deletion, represented as Diff2. Then, we defined the variation between single and double deletions by taking the average of Diff1 and Diff2 divided by the wild-type growth rate. The higher variation means that either the particular TF or metabolic gene is not essential for growth, while double deletion of this pair will decrease growth a lot. We identified 39 synthetic lethal or sick pairs of TFs and metabolic genes by setting the variation to greater than 90%. Moreover, most synthetic interacting pairs resulted in no growth, but single deletion of the corresponding TF or gene can maintain at least 95% of the wild-type growth.

We validated the predicted synthetic lethal or sick pairs by experimental growth assay. *Saccharomyces cerevisiae* single deletion strains were from the yeast deletion haploid collection (BY4742; Invitrogen). All double deletion strains were haploids generated by mating corresponding single deletion strains from the same library or from the BY4741 collection (Invitrogen), followed by tetrad dissection and selection by G418 resistance and PCR.

### ODELAY validation of synthetic lethal or sick pairs of TFs and metabolic genes

ODELAY was used to provide objective measurements of yeast growth defects (59). Yeast strains were cultured in YPD media in 96 well plates overnight. Cultures were diluted to an OD600 of 0.09 and allowed to grow for 6 hours at 30C. The cultures were then diluted to an OD600 of 0.02 and spotted onto YEP agarose media with 2% glucose. In ODELAY, colonies growing from individual cells are imaged and tracked using time-lapse microscopy for 48 hours with 30 minute intervals between images (59). All images were collected on Leica DMI6000 microscopes with a 10X 0.3NA lens using bright field microscopy. Colony area measurements are fit to the Gompertz function to estimate the colony doubling times. Between 100 and 300 cells growing into colonies were observed per strain. Estimated doubling times inform a confidence score identifying double deletion strains with synthetic growth defects (i.e. defects more severe than expected from the growth rates of constitutive single deletion strains). More details about the ODELAY analysis are available in the supplementary table (Figure S5) as well as the raw data (Table S7) and analysis script (Script S2).

## Acknowledgements

We thank Dr. Sriram Chandrasekaran and Dr. David Reiss for valuable discussions on the model integration. We gratefully acknowledge funding from the Shanghai Natural Science Funding / 16ZR1449700 (ZW), the Scientific Research Foundation for the Returned Overseas Chinese Scholars of the State Education Ministry / 15Z102050028 (ZW), the National Key Research and Development Plan of China / 2016YFC0902403 (ZW), the “SMC-SJTU Chen Xing” Program for Excellent Young Scholars / 13X100010027 (ZW), the open project from Synthetic Biology Key Laboratory of Chinese Academy of Sciences (ZW), an NSF Graduate Research Fellowship / DGE-1144245 (SM), a National Institutes of Health Center for Systems Biology / P50 GM076547 (JDA), the National Institutes of Health National Center for Dynamic Interactome Research / P41 GM109824 (JDA), U.S. National Science Foundation Awards MCB-1330912 and DB-1262637 (NSB), and a United States Department of Energy’s Advanced Research Projects Agency-Energy program / DE-AR0000426 (NDP).

## Supplemental files

**Figure S1**. Composition of the integrated models PROM and IDREAM.

A. The number of transcription factors in PROM and IDREAM. ‘Match_measuredTF’ is the number of TFs having a corresponding phenotype in Fendt’s experiment for 119 TF mutants. ‘Match_17defectTF’ is the number of TFs out of the 17 defect-inducing TFs that are involved in the two integrated models.
B. The log value of number of regulatory interactions and metabolic genes in PROM and IDREAM.

**Figure S2.** ROC curves for growth defect predictions with series of different thresholds using IDREAM and PROM on Yeast6 model. Across 16 different thresholds, the AUC value is significantly higher for IDREAM (mean = 0.67) than PROM (mean = 0.58).

**Figure S3**. ROC curve of IDREAM and PROM built on different yeast metabolic models (threshold=0.5). There are no significant differences by the three yeast models.

**Figure S4**. Predicted growth ratios for double deletions of TFs and metabolic genes using the IDREAM model. Each row represents a metabolic gene, and each column represents a gene encoding a TF.

**Figure S5**. The analysis of double knockout strain phenotypes by ODELAY. Estimated doubling times inform a confidence score identifying double deletion strains with synthetic growth defects. The first columns shows the metric and plate, such that ‘Mean.1’ means that for replicate 1, the average growth rate of many colonies was estimated using the mean and ‘Geometric Mean.2’ means that for replicate 2, the average growth rate was calculated using the geometric mean. ‘# Exp’ and ‘# Control’ refers to the number of strains in the experiment and control sets (respectively) where the growth rate in the double deletion was significantly less than that expected by adding together the growth rates for single deletions. ‘Mean Exp’ and ‘Mean Control’ are the mean growth decrease in growth rate beyond that expected from adding together the single deletion decreases in growth rate for the experimental and control sets (respectively). ‘Fisher p-Val’ uses a Fisher’s exact test to compare the ‘# Exp’ to the ‘# Control’ while ‘T-test p-Val’ uses the measured magnitudes of the synergistic growth defects.

**Table S1**. Flux variability for models with “frozen” state of fluxes constrained by each TF perturbation

**Table S2**. Significance test on MCC of IDREAM by randomly permuting the expression data and TF-gene interactions.

**Table S3**. Predicted growth ratio for TF knockouts using CLR-inferred regulatory network to link with the Yeast6 metabolic model.

**Table S4**. The comparison of AUC for growth defect predictions with series of different thresholds using IDREAM and PROM on Yeast6 model.

**Table S5**. Growth predictions with IDREAM by aggregate correlation.

**Table S6**. Yeast regulatory network from the YEASTRACT database and inferred by EGRIN.

**Table S7**. The raw doubling times for individual colonies as measured by ODELAY.

**Script S1.** The implementation of the IDREAM method.

**Script S2.** The R script for analysis of the results by ODELAY.

